## Supplementary Figures S1-S3 and Tables S1-S2 for "Unanticipated mechanisms of covalent inhibitor and synthetic ligand cobinding to PPARγ"

[1] Department of Integrative Structural and Computational Biology, Scripps Research and The Herbert Wertheim UF Scripps Institute for Biomedical Innovation & Technology, Jupiter, Florida 33458, United States
[2] School of Basic Medical Sciences, Guangzhou Laboratory, Guangzhou Medical University, Guangzhou, China.
[3] Department of Biochemistry, Vanderbilt University, Nashville, Tennessee 37232, United States.
[4] Center for Structural Biology, Vanderbilt University, Nashville, Tennessee 37232, United States
[5] Vanderbilt Institute of Chemical Biology, Vanderbilt University, Nashville, Tennessee 37232, United States
[6] Center for Applied AI in Protein Dynamics, Vanderbilt University, Nashville, Tennessee 37232, United States

File contains:

- Supplementary Figures S1–S3
- Supplementary Tables S1–S2

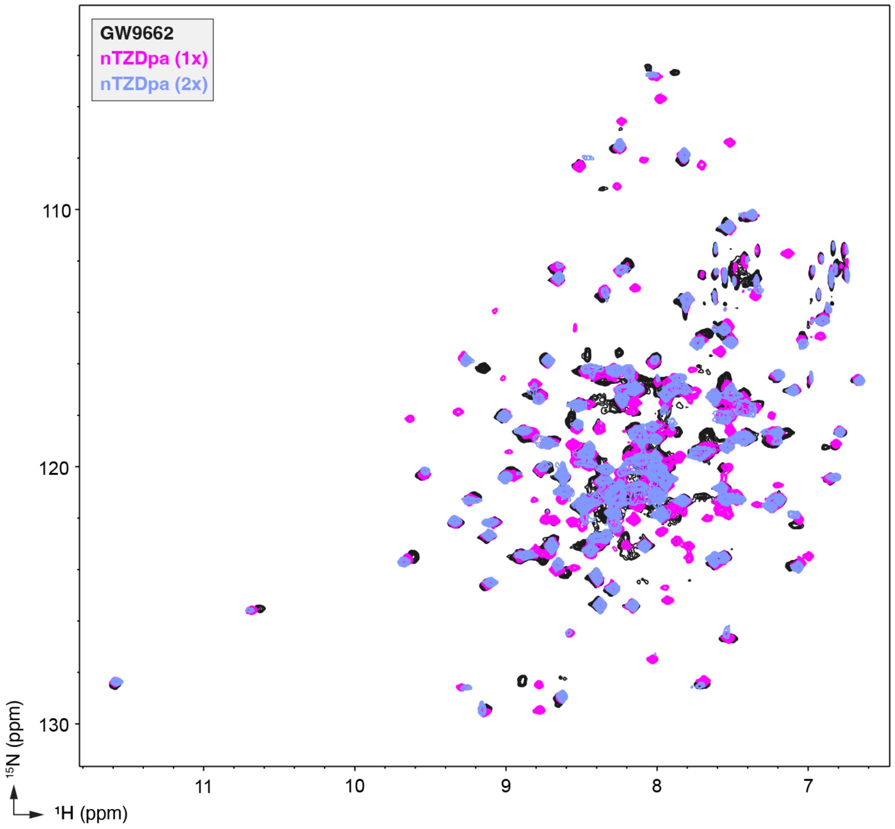


**Supplementary Figure S1. NMR analysis reveals PPARγ LBD binds more than one equivalent of nTZDpa.** 2D [$^{1}$H,$^{15}$N]-TROSY-HSQC NMR data of $^{15}$N-labeled PPARγ LBD in the absence or presence of nTZDpa added at the indicated molar equivalents.

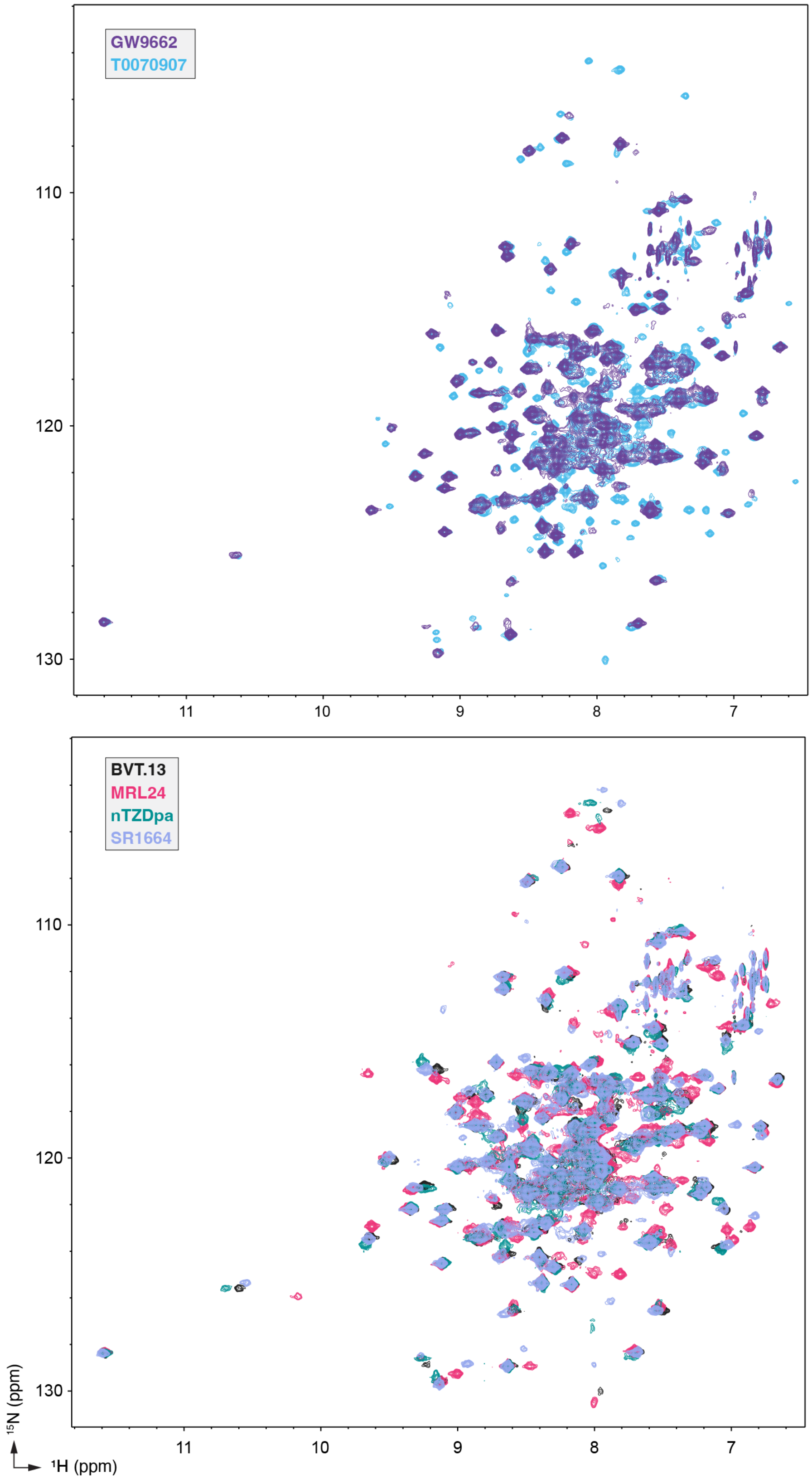


**Supplementary Figure S2. NMR spectral overlays show chemical shift perturbations (CSPs) between different single ligand-bound PPARγ LBD states.** 2D [$^1$H,$^{15}$N]-TROSY-HSQC NMR data of $^{15}$N-labeled PPARγ LBD in the presence of the indicated ligands added at 2 molar equivalents.

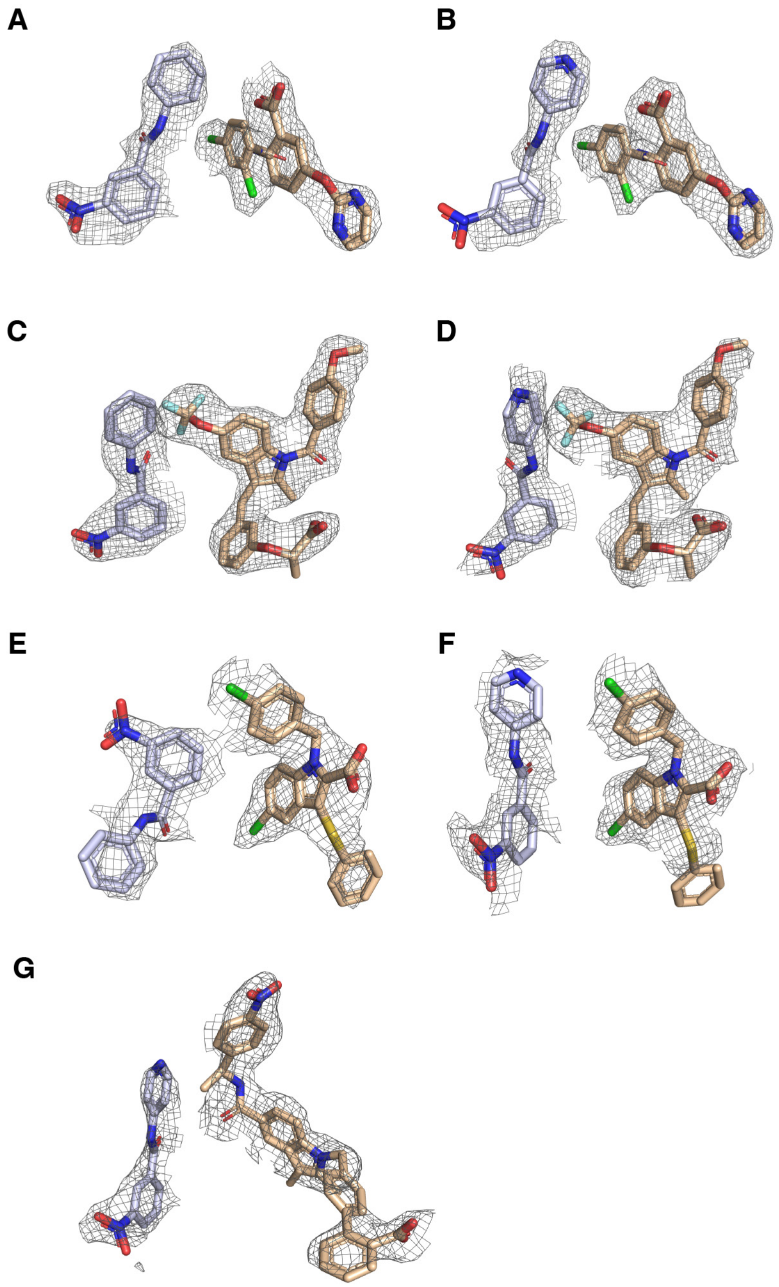


**Supplementary Figure S3. Electron density is shown from composite omit $2F_o$-$F_c$ maps (contoured at 0.8-1σ) of ligands for PPARγ LBD cobound to non-covalent ligands.** All densities are shown from chain B of the cobound structures for (**A**) GW9662 and BVT.13, (**B**) T0070907 and BVT.13, (**C**) GW9662 and MRL24, (**D**) T0070907 and MRL24, (**E**) GW9662 and nTZDpa, (**F**) T0070907 and nTZDpa, and (**G**) T0070907 and SR1664.

**Supplementary Table S1.** X-ray crystallography data collection and refinement statistics.

| | PPARγ LBD bound to GW9662 and BVT.13 | PPARγ LBD bound to GW9662 and MRL24 | PPARγ LBD bound to GW9662 and nTZDpa | PPARγ LBD bound to T0070907 and BVT.13 | PPARγ LBD bound to T0070907 and MRL24 | PPARγ LBD bound to T0070907 and nTZDpa | PPARγ LBD bound to T0070907 and SR1664 |
|---|---|---|---|---|---|---|---|
| **Data collection** | | | | | | | |
| Space group | C 1 2 1 | C 1 2 1 | C 1 2 1 | C 1 2 1 | C 1 2 1 | C 1 2 1 | C 1 2 1 |
| Cell dimensions | | | | | | | |
| *a*, *b*, *c* (Å) | 92.19, 61.99, 118.84 | 91.84, 62.22, 119.24 | 92.75, 62.22, 119.08 | 92.64, 61.78, 119.13 | 92.42, 61.63, 119.55 | 93.02, 62.16, 119.46 | 93.88, 62.68, 121.04 |
| α, β, γ (°) | 90, 102.38, 90 | 90, 102.28, 90 | 90, 102.19, 90 | 90, 102.38, 90 | 90, 102.19, 90 | 90, 102.14, 90 | 90, 102.46, 90 |
| Resolution (Å) | 51.06-2.54 (2.63-2.54) | 49.06-2.48 (2.57- 2.48) | 49.16-3.15 (3.26-3.15) | 51.02-2.49 (2.58-2.49 | 58.43-2.56 (2.65-2.56) | 58.39-2.73 (2.83-2.73) | 59.09-3.2 (3.31-3.2) |
| $R_{merge}$ | 0.088 (1.159) | 0.132 (1.791) | 0.043 (0.212) | 0.076 (1.083) | 0.087 (1.367) | 0.108 (1.621) | 0.046 (0.263) |
| $I / \sigma I$ | 12.06 (1.38) | 8.54 (1.04) | 15.02 (3.63) | 14.71 (1.76) | 12.80 (1.42) | 10.28 (1.21) | 10.70 (2.94) |
| Completeness (%) | 98.14 (96.97) | 98.41 (96.93) | 99.59 (100.00) | 99.14 (99.22) | 99.47 (98.55) | 98.79 (98.27) | 99.87 (100.00) |
| Redundancy | 6.6 (6.4) | 6.5 (6.5) | 2.0 (2.0) | 6.6 (6.6) | 6.6 (6.4) | 6.6 (6.7) | 2.0 (2.0) |
| **Refinement** | | | | | | | |
| Resolution (Å) | 2.54 | 2.48 | 3.15 | 2.49 | 2.56 | 2.73 | 3.2 |
| No. unique reflections | 21761 | 23546 | 11672 | 23269 | 21415 | 17965 | 11555 |
| $R_{work}$ / $R_{free}$ | 25.2/31.6 | 24.2/29.8 | 21.6/29.8 | 23.6/29.5 | 23.7/27.8 | 25.0/30.4 | 20.6/28.3 |
| No. atoms | | | | | | | |
| Protein | 4149 | 4065 | 3921 | 4123 | 4066 | 4096 | 4015 |
| Ligand/ion | 90 | 112 | 74 | 90 | 112 | 46 | 118 |
| Water | 35 | 82 | 2 | 43 | 21 | 18 | 0 |
| *B*-factors | | | | | | | |
| Protein | 65.80 | 53.94 | 58.48 | 64.00 | 64.71 | 67.70 | 79.58 |
| Ligand/ion | 74.57 | 41.56 | 77.08 | 74.58 | 49.66 | 83.90 | 108.68 |
| Water | 55.23 | 46.34 | 46.73 | 51.89 | 50.58 | 55.19 | n/a |
| R.m.s. deviations | | | | | | | |
| Bond lengths (Å) | 0.010 | 0.011 | 0.013 | 0.010 | 0.010 | 0.012 | 0.011 |
| Bond angles (°) | 1.33 | 1.45 | 1.43 | 1.21 | 1.38 | 1.35 | 1.31 |
| Ramachandran favored (%) | 92.90 | 96.77 | 94.33 | 96.63 | 96.57 | 95.01 | 92.61 |
| Ramachandran outliers (%) | 0.20 | 0.20 | 0.00 | 0.00 | 0.20 | 0.20 | 0.21 |
| PDB accession code | 8ZFN | 8ZFP | 8ZFO | 8ZFQ | 8ZFS | 8ZFR | 8ZFT |

*Values in parentheses are for highest-resolution shell.

**Supplementary Table S2.** Structural rmsd comparison of ligand cobound structures to the transcriptionally active PPARγ LBD conformation (PDB 6ONJ).

| **PDB ID** | **rmsd** |
|---|---|
| 6ONJ | n.d. |
| 8ZFP | 0.98 |
| 8ZFO | 0.77 |
| 8ZFQ | 0.97 |
| 8ZFS | 0.95 |
| 8ZFR | 0.92 |
| 8ZFT | 1.00 |
| 8ZFN | 1.03 |
